# Understanding consumers to inform market interventions for Singapore’s shark fin trade

**DOI:** 10.1101/2023.06.27.545387

**Authors:** Christina Choy, Hollie Booth, Diogo Veríssimo

**Author notes:** Corresponding author: Hollie Booth.

## Abstract

1. Sharks, rays and their cartilaginous relatives (Class Chondricthyes, herein ‘sharks’) are amongst the world’s most threatened species groups, primarily due to overfishing, which in turn is driven by complex market forces including demand for fins. Understanding the high-value shark fin market is a global priority for conserving shark and rays, yet the preferences of shark fin consumers are not well understood. This gap hinders the design of evidence-based consumer-focused conservation interventions.
2. Using an online discrete choice experiment, we explored preferences for price, quality, size, menu types (as a proxy for exclusivity) and source of fins (with varying degrees of sustainability) among 300 shark fin consumers in Singapore: a global entrepot for shark fin trade.
3. Overall, consumers preferred lower-priced fins sourced from responsible fisheries or produced using novel lab-cultured techniques. We also identified four consumer segments, each with distinct psychographic characteristics and consumption behaviors.
4. These preferences and profiles could be leveraged to inform new regulatory and market-based interventions regarding the sale and consumption of shark fins, and incentivize responsible fisheries and lab-cultured innovation for delivering conservation and sustainability goals.
5. In addition, message framing around health benefits, shark endangerment and counterfeiting could reinforce existing beliefs amongst consumers in Singapore and drive behavioral shifts to ensure that market demand remains within the limits of sustainable supply.

## 1. Introduction

Overexploitation threatens the survival of wild populations and puts species at risk of extinction (Maxwell et al., 2016). More than one third of sharks, rays, and chimera (Class Chondrichthyes, herein sharks), especially those which are large-bodied and living in shallow waters, are highly threatened primarily due to overfishing. This makes them one of the most threatened groups of vertebrates (Dulvy et al., 2014; Mull et al., 2022). Oceanic populations have declined by over 70% in the past 50 years, and reef sharks are functionally absent from nearly 20% of the world’s coral reefs (MacNeil et al., 2020). Moreover, the slow life history traits of many shark species (slow growth, late age of maturity, low fecundity, and long gestation period) make it especially difficult for their populations to withstand fishing pressure and recover from historic overfishing (Cortés, 2002; García, Lucifora & Myers, 2008; Dulvy & Forrest, 2010; Dulvy et al., 2014). Yet as apex- and meso-predators sharks play vital roles in maintaining healthy and functional marine ecosystems (Stevens et al., 2005; Ferretti et al., 2010; Heithaus et al., 2012), and generate economic and subsistence value via marine tourism and the fisheries livelihoods of many coastal communities (Gallagher & Hammerschlag, 2011; Booth, Squires & Milner-Gulland, 2019; Mustika, Ichsan & Booth, 2020). As such, conservation actions are urgently needed to halt population declines, prevent extinctions and promote recovery of shark populations and marine ecosystems (Dulvy et al., 2014).

Sharks are caught throughout the world’s fisheries as both target and incidentally-caught species; and retained and sold for their cartilage, meat, skin, liver oil and fins (Dulvy et al., 2014; Oliver et al., 2015). Notably, the fins of certain species have particularly high value, driven largely by demand for shark fin soup, a popular delicacy among many Asian communities across the globe (Choy & Wainwright, 2022). Shark fin soup is made primarily from the ceratotrichia (fibrous cartilage) within shark fins. It has been traditionally served as a signal of power in Chinese folk custom, and is believed to provide tonic health benefits to consumers (Clarke, Milner-Gulland & BjØrndal, 2007; Fabinyi, 2012; Dell’Apa, Chad Smith & Kaneshiro-Pineiro, 2014). This dish is usually served at important corporate or social banquets as a prestigious item believed to embody wealth, hospitality, and social status; and thus reinforce relationships among people (Dell’Apa, Chad Smith & Kaneshiro-Pineiro, 2014; Zhou et al., 2021). One study in China found that consumers generally prefer higher priced and rarer shark-fins, which reinforces the notion that shark fin is consumed as a luxury goods (Zhou et al., 2021). As such, fins are often sold at US$200-400 per kilogram, and sometimes up US$1,000 per kilogram (Clarke, 2004; Southeast Asian Fisheries Development Center, 2006).

In total, it is estimated that the fins of between 26 and 73 million sharks, worth US$400-550 million, are traded each year globally (Clarke, Milner-Gulland & BjØrndal, 2007). In turn, this high value market creates an incentive for fisheries to target sharks or retain incidentally caught sharks. Since the fins are more valuable than the rest of the body, shark ‘finning’ also sometimes occurs, wherein fins are retained and carcasses are discarded at sea (Dell’Apa, Chad Smith & Kaneshiro-Pineiro, 2014). Overall it is estimated that almost 94% of sharks currently in the fin trade are derived from unmanaged sources (Oliver et al., 2015). While a handful of bright spots of good shark fisheries management do exist (Simpfendorfer & Dulvy, 2017), these typically do not occur in hotspots of conservation need and fishing threat (Dulvy et al., 2017). Such hotspots typically occur in less economically developed countries, which are often dominated by small-scale fisheries, and where good management is hindered by socio-economic complexities, and a lack of data and capacity (Booth, Squires & Milner-Gulland, 2019). Moreover, though there have been recent attempts to improve global management of shark populations, e.g., through regulating fishing and trade of sharks via fin bans and species-specific trade controls, there remains limited evidence that these policies have led to widespread positive conservation outcomes for sharks (Clarke et al., 2013; Worm et al., 2013; Dulvy et al., 2014). This may be due to complexities surrounding the micro- and macro-economic drivers of shark fishing, which cannot simply be addressed through direct regulation (Booth, Squires & Milner-Gulland, 2019). As such, more nuanced and concerted efforts are still needed, with novel regulatory and market-based reforms to drive transformative changes in fisheries management.

Within this global context, Asia is a priority region for shark conservation, since it is home to hotspots of species diversity and endemicity, hotspots of fishing pressure, and hotspots of consumer demand (Dulvy et al., 2017). Of countries in this region, Singapore is among the top importers/re-exporters and consumer countries of shark fins, where critically endangered hammerhead sharks and wedgefishes are amongst the species regularly sold and consumed (Wainwright et al., 2018; Choo et al., 2021; Liu et al., 2021; Choy & Wainwright, 2022). While several food establishments have recently removed shark fin soup and other shark products from their menus, trade data between 2017 and 2020 implies that local appetite for shark fins persists (Yeo, 2022). Creating a more sustainable pathway for shark fin consumption in Singapore (e.g., via demand reduction and/or sustainable use) will require nuanced and culturally relevant interventions. Yet the nature of demand for shark fins in Singapore remains poorly understood. Aside from basic information on prices and sales (Choy et al., 2022), there is little-to-no information on consumer demographics and consumption patterns.

In addition, Singapore offers an interesting context to understand consumer preferences for lab-cultured shark fin as a sustainable alternative, since it is a leading market for lab-cultured protein sources (Phua, 2020; Chriki, Ellies-Oury & Hocquette, 2022). Shark fins have been developed using lab-cultured techniques and are currently available in the retail market at a price more affordable than wild-sourced shark fins (Alpha Food Labs, 2022). These novel food technologies offer major opportunities to augment supply, meet existing demand for fins, and alleviate harvest pressure on wild populations. However, commercial and conservation success of lab-cultured alternatives heavily depends on whether consumers are willing to try and buy them. Previous studies have reported consumer concerns surrounding food safety; nutritional value; and the taste, texture and appearance of lab-cultured meat (Mancini & Antonioli, 2020). In addition, since people with different demographics and cultural backgrounds have varying attitudes towards cultured meat (Zhang et al., 2021), it is necessary to understand the extent to which shark fin consumers in the Singapore market may prefer lab-cultured alternatives over wild-sourced shark fins.

To fill this gap, we conducted a Discrete Choice Experiment (DCE) with shark fin consumers in Singapore that aimed to (a) identify consumer segments with distinct preferences for shark fin soup, based on socio-demographic characteristics; and (b) understand the drivers and circumstances for consuming shark fin soup. This study is also among the first to assess substitutability of wild shark fins with lab-cultured alternatives, and thus could inform innovative market-based approaches to reduce fishing pressure on endangered species.

## 2. Methods

### 2.1 Survey questionnaire design

The survey questionnaire was designed in English, the official and commonly spoken language in Singapore, with four parts (Supplemental Material). Part 1 (Questions 1 to 6) obtained respondents’ demographic characteristics including age, gender, race, religion, education level, monthly income range, and if applicable, their Chinese dialect (Doughty et al., 2021a). Part 2 (Questions 7 – 13) explored respondents’ purchasing and consumption patterns of shark fin in the past year, to gauge the frequency of recent consumption, purchase location, and trends in consumption. Part 3 (Questions 14 and 15) assessed the attitudes of respondents regarding consuming shark fins and the ecological importance of sharks, using a 5-point Likert scale. We also used the Schultz scale to measure a respondent’s environmental concern, where respondents are asked to rate nine items from three dimensions representing biosphere concern (plants, marine life, birds, animals), egoistic concern (me, my health, my future) and altruistic concern (all people, children) (Schultz, 2001; Cruz & Manata, 2020). This scale was chosen over other instruments for its reliability, brevity, and popularity among other studies which makes comparisons across different sample populations possible (Cruz & Manata, 2020). The final section was the Discrete Choice Experiment (DCE), which was designed based on a shark fin soup consumption scenario, where each soup alternative had five varying attributes: price, size, texture, source of fins and type of menu (Figure 1).

**Figure 1.**
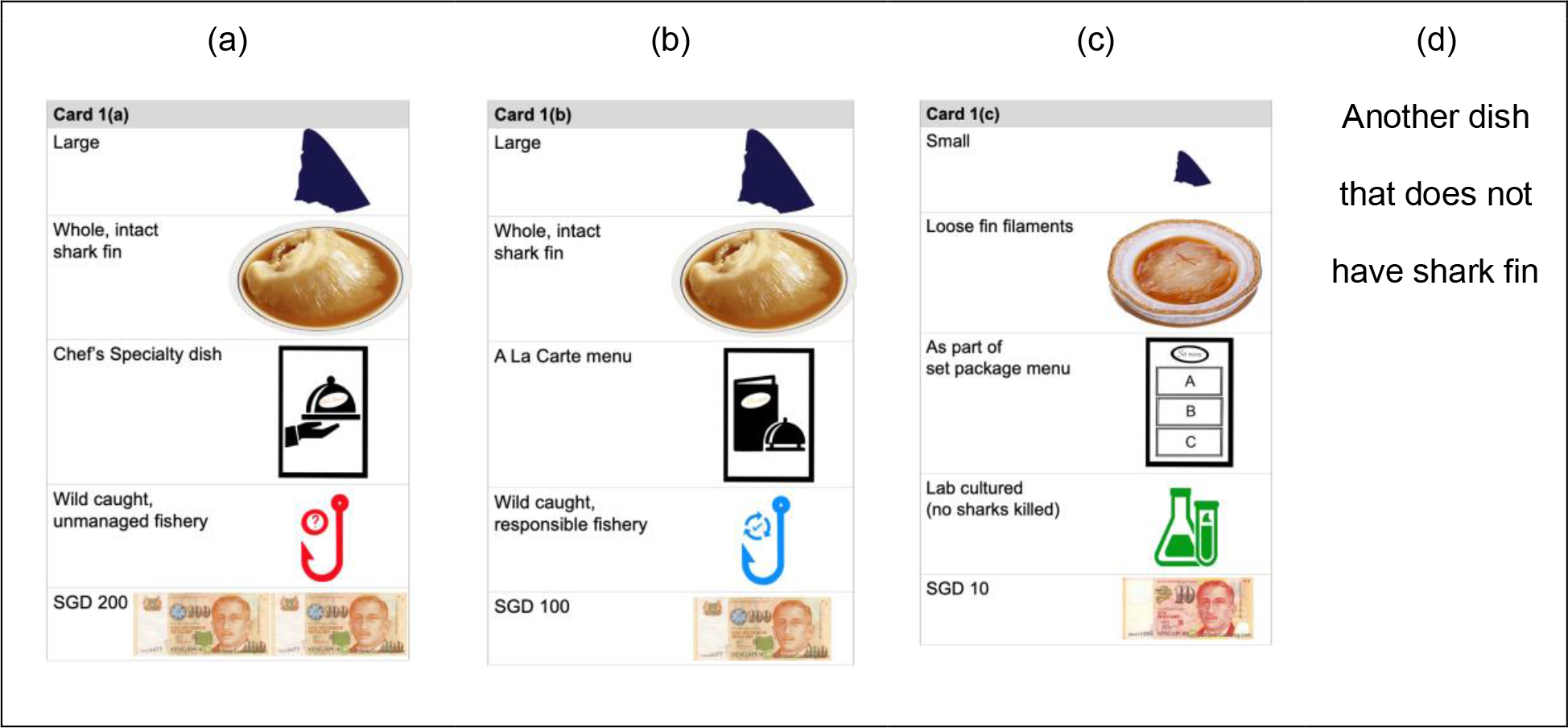
Sample set of choice cards (i.e., alternatives) with different levels for attributes, used for the discrete choice experiment on shark fin consumption in Singapore.

### 2.2 Discrete Choice Experiment design

We designed the DCE based on an initial scoping phase, which included a literature review and informal discussions with key stakeholders (i.e., restaurants, dried seafood retail shops), to ensure the scenario and attributes were relevant and reflected real purchasing decisions.

To ensure that respondents understood the circumstances under which they are required to make a choice, we first presented them with a short paragraph detailing the scenario of purchasing shark fin soup in a Chinese restaurant when dining with their family (including older folks). This scenario is thought to be common, based on information from the scoping phase. A “cheap talk script” was also included to remind respondents to behave as if real transactions were taking place, to mitigate “hypothetical bias” (i.e., that responses in our hypothetical valuation differ from choices in real life) (Tonsor & Shupp, 2011; Doughty et al., 2021a).

Respondents were presented with four alternatives, which included three shark fin soup options with different combinations of attributes (Zhou et al., 2021), and a “none” alternative to offer a realistic scenario in which respondents could choose to opt out if none of the shark fin alternatives were satisfactory (Figure 1). This is in line with consumer theory, and thus preferred despite the loss of efficiency (Doughty et al., 2021a). Each alternative consisted of five attributes with differing levels, as follows: Price is a typical attribute used in DCEs to measure willingness to pay. A total of five levels, ranging between SGD $10 and SGD $200 were used for the price, reflecting the actual cost per bowl of shark fin soup offered in restaurants in Singapore (based on scoping data). Both size and quality of fins were included as attributes, each with two levels (Table 1) where larger and more intact fins are typically perceived as higher quality based on scoping data. The source of the fin was included as a conservation- or management-relevant attribute, since previous research with consumers in China indicated willingness-to-pay for responsibly-sourced fins (Zhou et al., 2021) and our study sought to understand preferences for conservation-friendly sources including lab-cultured alternatives. Since consumers can also purchase shark fin soups ala-carte, as part of set menu alongside with other dishes, or as a specialty dish at restaurants in Singapore, it is likely that the different marketing strategies and business models could affect consumer choices; and that these different choices may reflect different perceived levels of luxury or prestige (Table 1). As such we included three different types of menus as a proxy for perceived luxury or specialty. Based on this design, respondents were then asked to select which shark fin soup they would purchase based on the soups available in each round. Based on Table 1, we built a pilot orthogonal design using IBM SPSS 27.0 with the initial choice alternatives being paired using a “shifted technique” (Louviere, Hensher & Swait, 2000) into 25 dichotomous choices. The pilot survey received 21 responses between January and February 2022 using the SmartSurvey online survey platform (SmartSurvey, 2022). Data collected from the pilot survey were analyzed using a Multinomial Logit (MNL) model in NLOGITVersion 6.0. The results were used to obtain Bayesian priors to generate a D-efficient Bayesian design in Ngene 1.0.1 for the final DCE design (ChoiceMetrics, 2012). The final design contained 15 choices and a d-error of 0.134, the lowest across 100,000 iterations. Question ordering was randomized to mitigate the effects of respondent fatigue on results.

**Table 1.** Attributes and levels of a discrete choice experiment on shark fin soup consumption in China

| Attribute | Levels | Rationale for attribute and levels | Hypothesized preference |
| --- | --- | --- | --- |
| Price | SGD \$10<br>SGD \$50<br>SGD \$100<br>SGD \$150<br>SGD \$200 | Price range is based on cost of shark fin soup bowl (person serving) from actual restaurants. | Higher price - as consumers in China showed preference for expensive fins |
| Size of fin | Small<br>Large | Size of fin is based on whether a large species (e.g., wedgefish), individual size or certain part of the shark is preferred, which has implications on the species' wild stocks and status. | Larger fin - as it looks more presentable and luxurious |
| Texture and quality of fin | Loose shark fin filaments<br>Whole, intact shark fin | The filaments and thickness of the fins are an indicator of its authenticity and grade. | Whole, intact shark fin - as it looks more presentable and luxurious |
| Types of menus | As part of set package menu<br>A La Carte menu<br>Chef's Specialty dish | Types of menus are based on the different menus which consumers can purchase shark fin soup bowl from actual restaurants, which hints towards the exclusivity of the dish. | Chef's Specialty dish – as it may be associated with more prestige or luxury |
| Source of fin | Wild caught, unmanaged fishery<br>Wild caught, responsible fishery<br>Lab cultured (no sharks killed) |  | Unknown – to be tested in this study |

### 2.3 Data collection

We worked with Rakuten Insight Singapore (Rakuten Insight, 2022a) to disseminate the survey questionnaire via their in-house online panel between March 23 and April 4, 2022. Participants were pre-recruited and gave their consent to take part in market research studies. With the low incidence rate in Singapore for shark fin consumption, having a readily available group of individuals presented many advantages for our study, including the ability to reach specific audiences, rapid turnaround, higher participation rates and cost-effectiveness. Participants in Rakuten Insight’s panels earn points from online surveys which can be exchanged for gift cards and rewards such as vouchers and promotion codes. The online surveys can be taken from participants’ smartphone or desktop computer during their free time (Rakuten Insight, 2022b).

Of the 1,075 people who accessed the survey, 634 were screened out based on eligibility criteria, primarily due to them being non-consumers of shark fin. Of those eligible for the survey questionnaire, 88 did not complete the survey, giving a completion rate of 80%. A total of 353 completed responses were collected, however 53 responses were disqualified based on quality control checks (e.g., speed, duplicate answers, nonsense open-ended answers, straight answer patterns, logic check for inconsistent answers), giving a final sample of 300 individuals who have consumed shark fins and resided in Singapore over the past three years. A summary of the survey dispositions (screen outs, completes and incompletes) is provided in the Supplemental Material, Figure S1.

This study received ethical review and approval from the Wildlife Conservation Society’s Institutional Review Board (REF# 21-65).

### 2.4 Data analysis

Scale scores were produced for biospheric, egoistic, and altruistic concerns of each respondent by averaging the items in each domain (i.e., the items for biospheric are ‘plants’, ‘marine life’, ‘birds’, ‘animals’; for egoistic concerns are ‘me’, ‘my health’, ‘my future’; for altruistic concerns are ‘all people’, ‘children’) (Schultz, 2000).

To analyse the DCE, we first used a Multinomial Logit (MNL) model to assess the aggregate preferences of respondents. Next, we used a Latent Class Model (LCM) on LIMDEP NLOGIT 4.0 to explore potential heterogeneity in preferences and to partition the sampled population into more homogeneous classes. LCMs assume that respondents can be divided into a finite number of latent classes (or segments), based on their preferences, and then aim to characterize each of these segments based on observable respondent characteristics, most commonly socio-demographic variables (e.g., gender, age, monthly income, education, race). It should be noted that this characterisation of respondents is done in relation to a reference group and not in absolute terms.

We selected the model with the optimal numbers of consumer segments which is the most parsimonious by all three statistical criteria [e.g., Akaike Information Criteria (AIC) and Bayesian Information Criteria (BIC)]. The model was also determined by the size of a class membership, where classes with very small segments that have a higher probability of being spurious are avoided. We included an Alternative Specific Constant (ASC) to account for the “neither” responses. In all our analysis, the ASC took a value of 1 when “neither” choice was opted, reflecting the utility derived from not choosing any of the offered choice options.

## 3. Results

### 3.1 Demographic characteristics of sample population

Respondents were distributed across five age categories, and males comprised 60% of the sample population. Most respondents were Chinese (76%), and 38% of the Chinese respondents identified themselves as Hokkien. Christianity was the most represented religion (33%). More than half of the sample population have attained university (51%) and higher educational qualifications (20%), and 40% of the respondents have a monthly income of Singapore Dollars (SGD) $3,000 - $7,000. A summary of respondent demographics is available in Supplemental Material, Table S2.

### 3.2 Purchase and consumption habits and attitudes towards consuming shark fins

Respondents who reported having consumed shark fins in the past year comprised 33% of the sample population, closely followed by consumption within the past month (25%). A third of the sample population indicated consuming shark fins a few times per year. Respondents commonly consumed shark fins as part of weddings or special celebratory events, and thus did not purchase it themselves. A large proportion of respondents (61%) reported a decrease in their consumption in the past five years. However, there was no change in consumption in the past five years for 26% of the respondents, while 13% of respondents reported a large or small increase in their consumption within the past five years. A summary of the respondents’ purchase and consumption habits, as well as reasons for increased consumption, is available in Supplemental Material, Table S3.

In terms of attitudes toward consuming shark fins, there was wide agreement across respondents that shark fins are delicious (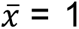 = 1, agree = 46%, strongly agree = 31%), sharks are important for the functioning of marine ecosystems (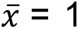 = 1, agree = 52%, strongly agree = 27%) and we should manage shark populations to sustain fish stocks (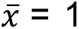 = 1, agree = 49%, strongly agree = 21%) (Figure 2; Supplemental Material, Figure S4 and Figure S5).

**Figure 2(a).**
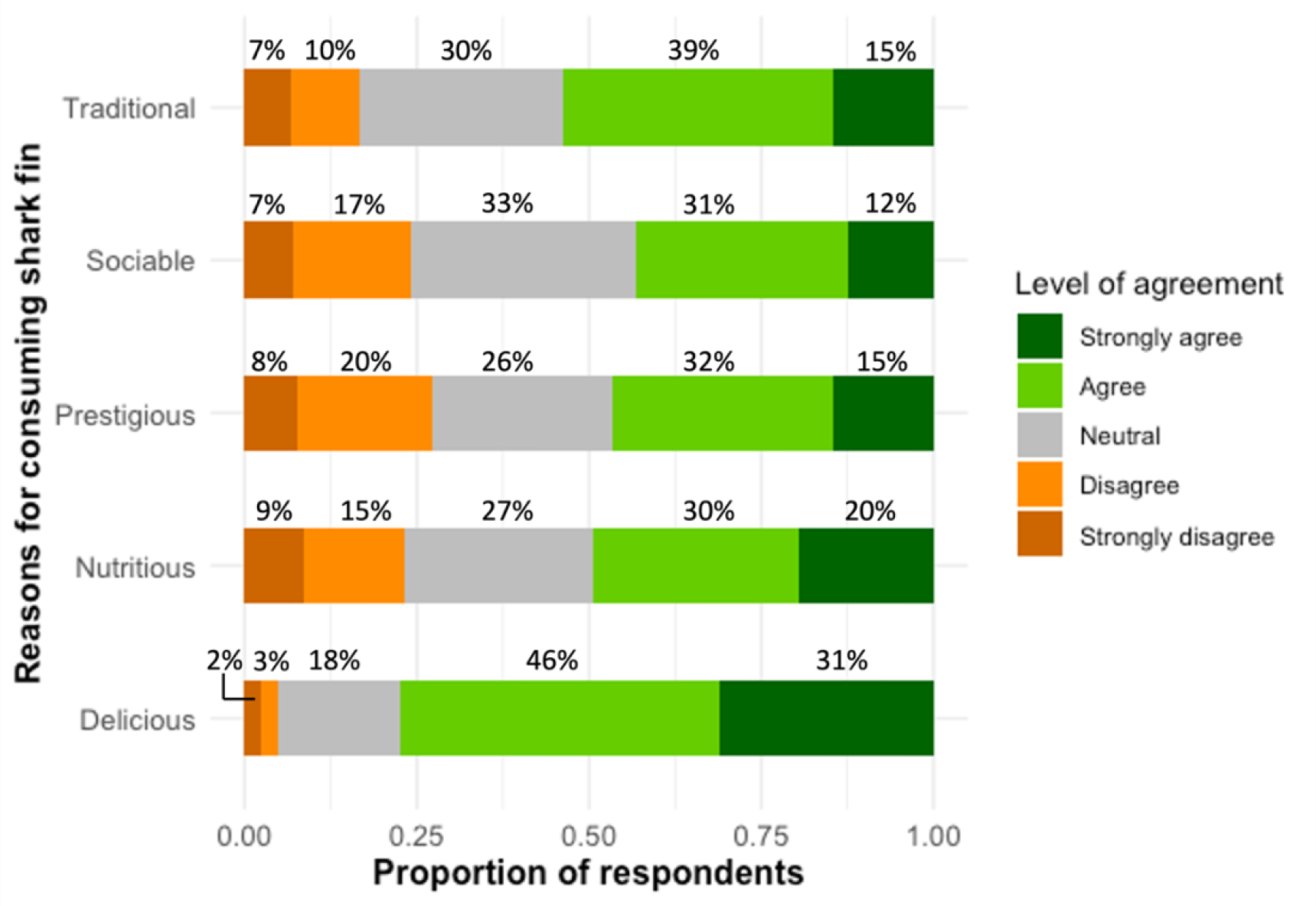
Proportion of respondents on reasons for consuming shark fins

**Figure 2(b).**
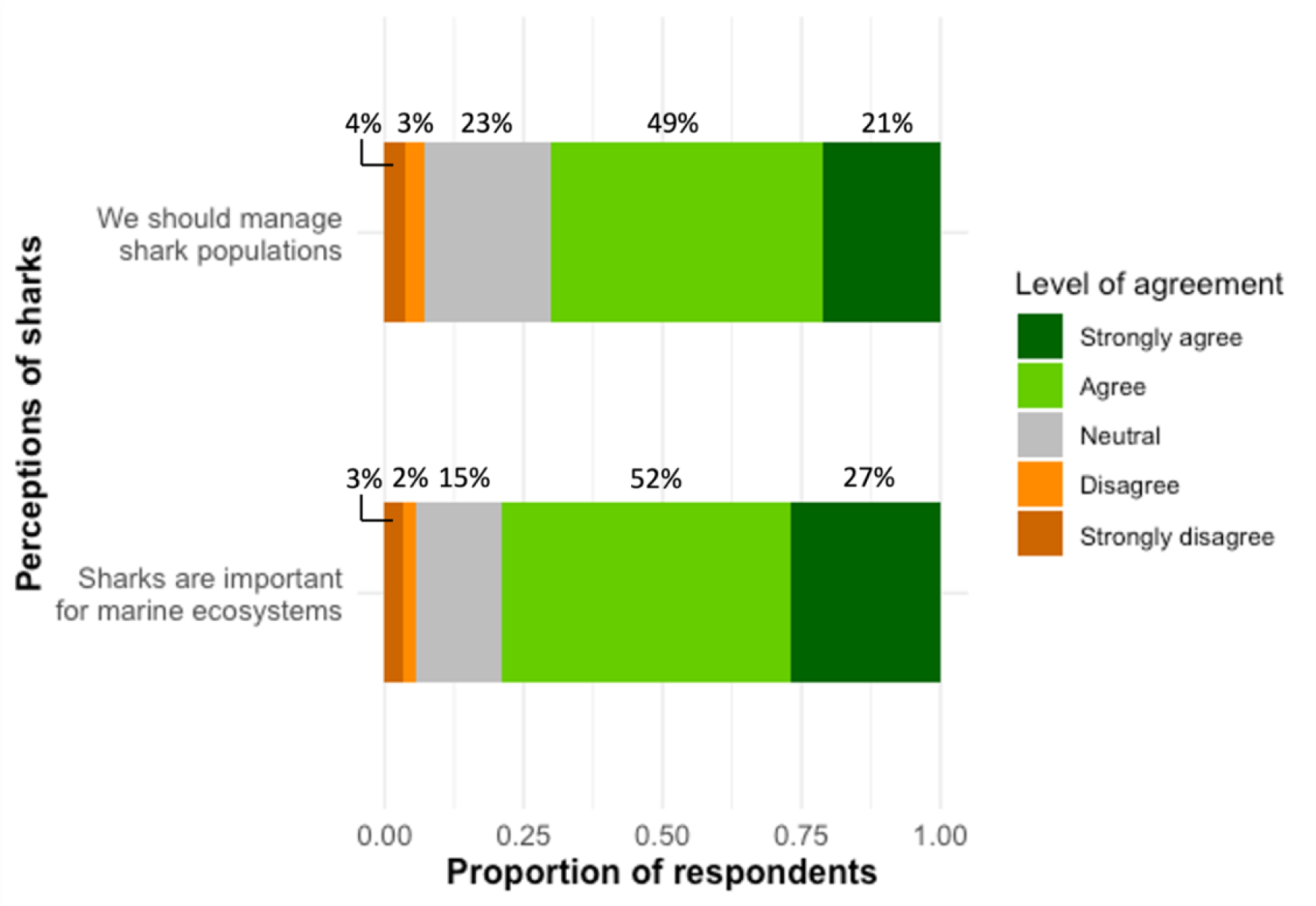
Proportion of respondents on perceptions of sharks

### 3.3 Choice experiment

To understand which factors were most relevant for consumer segmentation, we explored several LCM model specifications related to both respondent demographics and psychographics. The best performing LCM was segmented based on consumption frequency (Q8), consumption over the past five years (Q12), passive consumption (Q10) and egoistic environmental concern (Q15) (Table 2 and Supplemental Material, Survey questionnaire). The LCM model with five respondent segments was the most statistically efficient, however it performed poorly in several ways. For instance, it had two very small segments with less than 10% of the sample population in each segment, which could be dictated by the idiosyncrasies of a few participants in the sample population and therefore does not inform broader consumer trends. We therefore chose the model with four segments to explain heterogeneity across the sample population. This model was the second most parsimonious after the five-segment model, by all three statistical criteria in our analysis (Supplemental Material, Table S6).

**Table 2.**
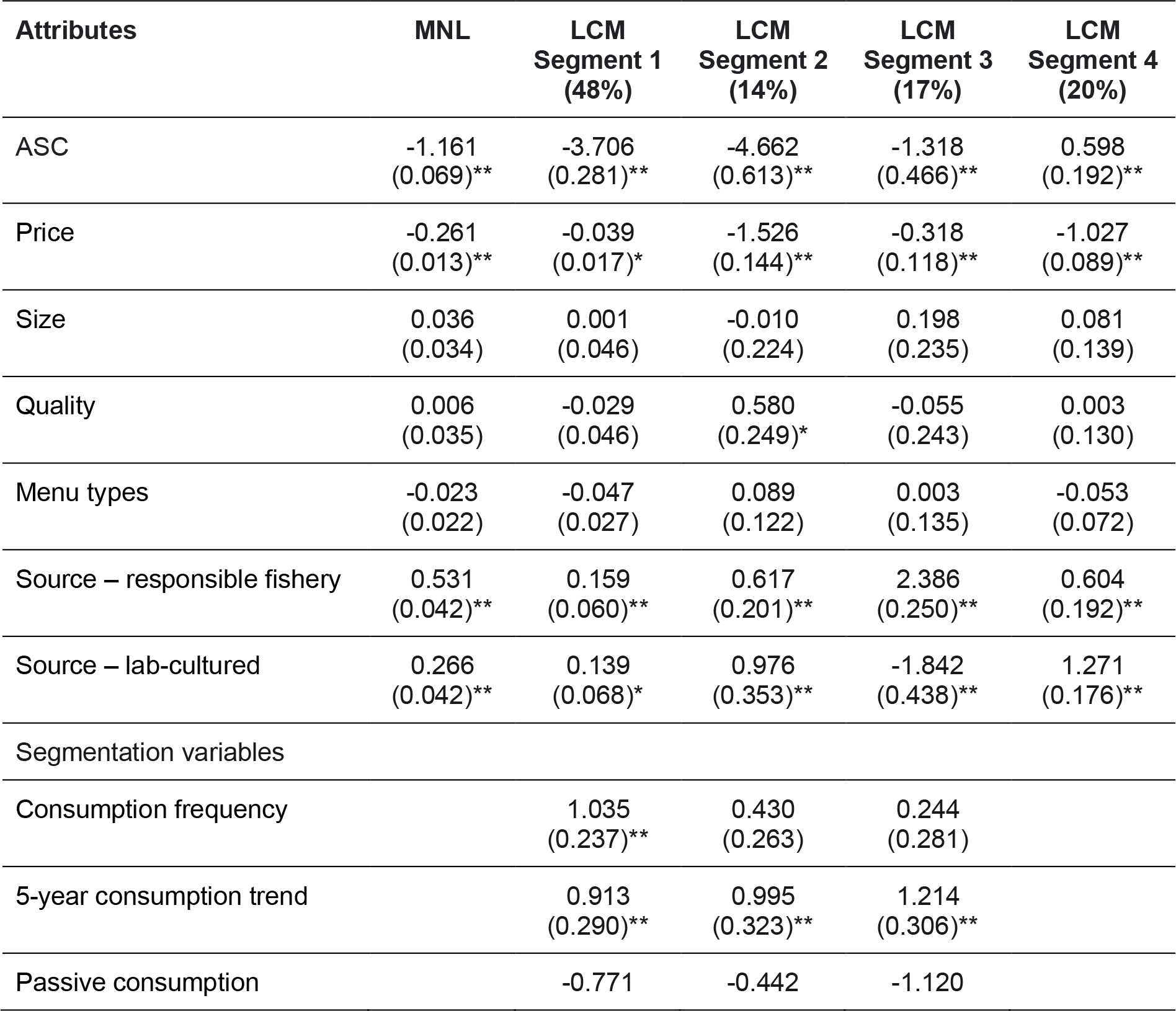
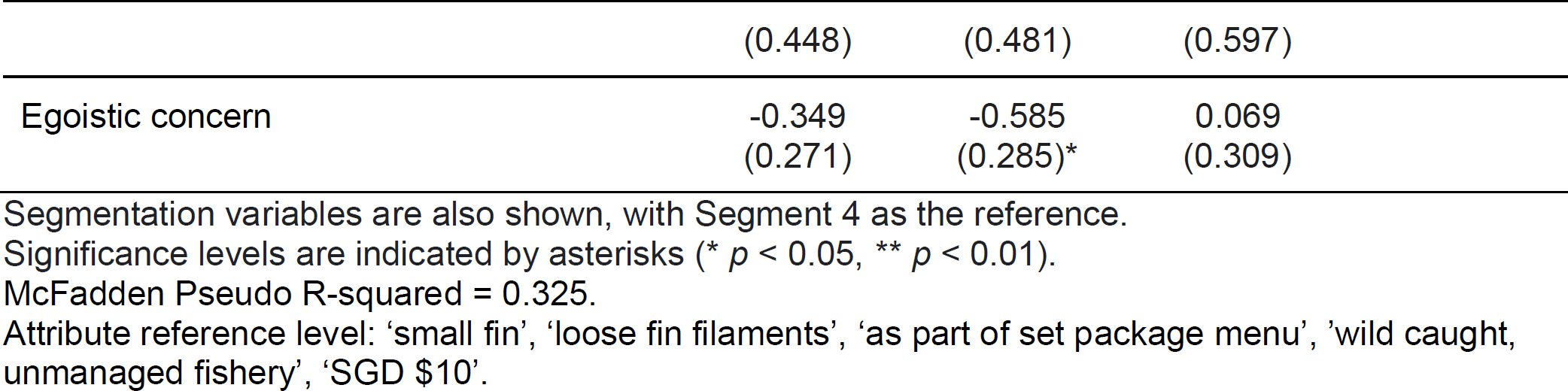
Multinomial Logit (MNL) and Latent Class Model (LCM) estimates of utility function for each attribute, with standard errors in parentheses (95% confidence intervals).

Both price and source of fins were statistically significant across the MNL and LCM estimates, indicating a preference for lower-priced and more sustainable (responsible fishery or lab-cultured) options. Size and quality of fins as well as the different menu types were not considered important overall (Table 2). We also calculated the willingness to pay for different attributes (Table 3). As the characterisation of respondents is done in relation to a reference group and not in absolute terms, the positive coefficients of the LCM estimates for consumption frequency and 5-year consumption trend across LCM Segments 1, 2 and 3 suggested that the segment of reference (Segment 4) consists of the minority who have low consumption frequency and reduced consumption a lot over the past five years.

**Table 3.**
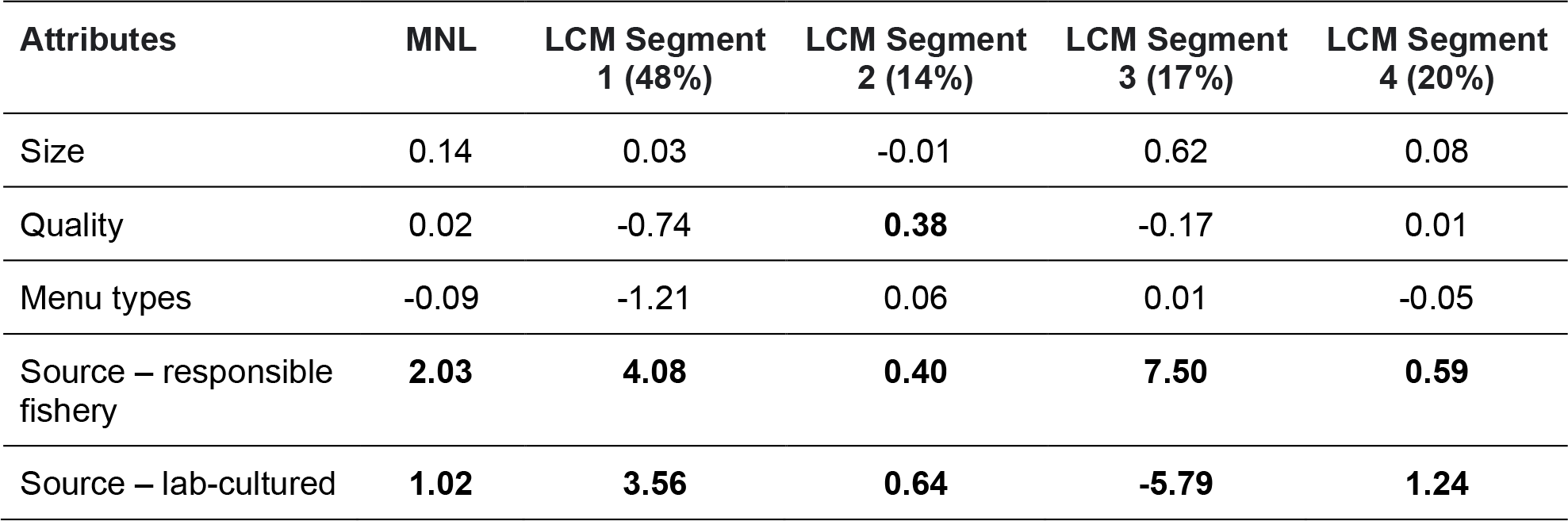
. Willingness to pay for different attributes of shark fin in Singapore. All values are in Singapore Dollars. Bold values are those relating to attributes that are statistically significant in terms of influencing choice.

| Attributes | MNL | LCM Segment<br>1 (48%) | LCM Segment<br>2 (14%) | LCM Segment<br>3 (17%) | LCM Segment<br>4 (20%) |
| --- | --- | --- | --- | --- | --- |
| Size | 0.14 | 0.03 | -0.01 | 0.62 | 0.08 |
| Quality | 0.02 | -0.74 | **0.38** | -0.17 | 0.01 |
| Menu types | -0.09 | -1.21 | 0.06 | 0.01 | -0.05 |
| Source – responsible fishery | **2.03** | **4.08** | **0.40** | **7.50** | **0.59** |
| Source – lab-cultured | **1.02** | **3.56** | **0.64** | **-5.79** | **1.24** |

Overall, there is a substantial homogeneity across the four segments, where Segment 1 is the largest and comprises almost half of the sample population who preferred sustainably sourced (from responsible fisheries or lab-cultured) and lower-priced fins. Members of Segment 1 had a significantly higher frequency of shark fin consumption, and had increased their consumption over the past five years, relative to Segment 4.

Like Segment 1, Segment 2 also preferred sustainably sourced and lower-priced fins but stood out for preferring higher quality fins. However, consumers in Segment 2 differed from Segment 1 by having significantly lower egoistic environmental concern. Segment 3 also comprised members who favored shark fins from responsible fisheries, but disliked lab-cultured shark fins. Consumers in this segment had significantly increased their consumption over the past five years, much more than the three other consumer segments. The level of egoistic environmental concern was also higher in this segment of consumers.

In contrast to Segment 3, members in both Segments 2 and 4 showed a strong preference for lab-cultured shark fins over fins obtained from unmanaged fisheries, suggesting that both segments could be early-adopters of sustainability innovations. Lastly, consumers in all four segments showed a preference for the “none” alternative and were primarily passively consuming shark fins. Figure 3 provides a summary of characteristics, preferences, and interventions for the four identified consumer segments.

**Figure 3.**
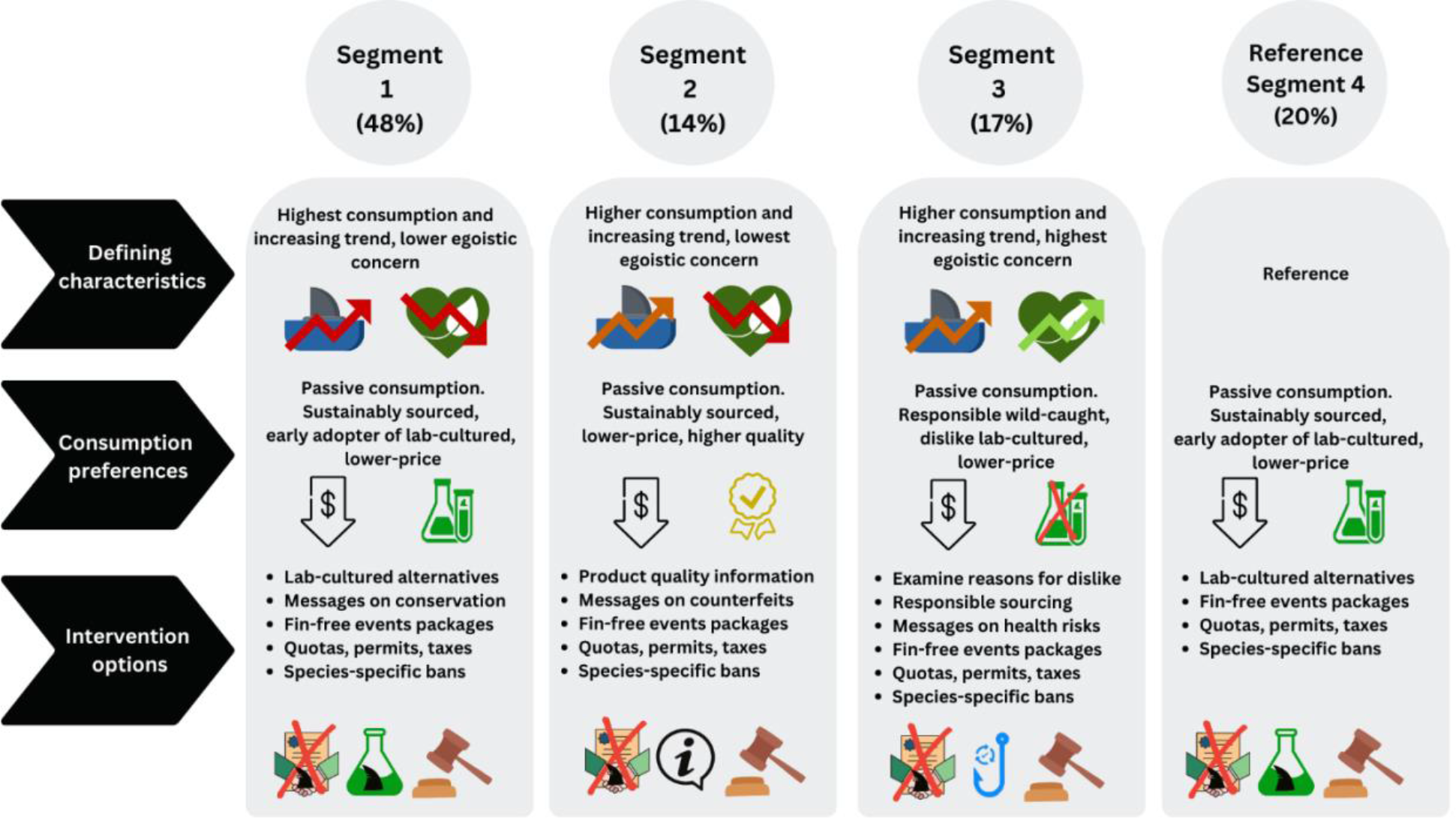
Characteristics, preferences, and interventions for the four identified consumer segments

## 4. Discussion

This study is the first to apply a discrete choice experiment to understand the demographics and preferences of shark fin consumers in Singapore, as well as the relative acceptance of lab-cultured alternatives to wild-sourced shark fins. Promisingly, we found that all respondents exhibited preferences for more responsibly and ethically sourced shark fins. As expected, price was also a significant factor which influenced consumer choices between potentially substitutable products, which validates our findings. We also identified four distinct segments of consumers, each with slight differences in their demographics and consumption behaviors, which has implications for future consumer-focused interventions.

### 4.1 Interpreting consumer preferences

#### 4.1.1 Preferences for lower-priced fins

Our findings revealed a negative relationship for price across all segments, indicating a preference for lower-priced shark fins. This preference is intuitive for most consumer goods. However, price is often used by consumers to judge a product’s quality, authenticity or luxury; and for luxury goods such as shark fins, other studies have documented that consumers are more likely to favor higher prices, as a signal of wealth and social status and fear of consuming counterfeit products (Zhou et al., 2021). However, this form of ‘conspicuous consumption’ appears to be less important for consumers in Singapore (Postrel, 2008; Stewart, 2022). There is a growing affluent population in Singapore, and recent studies have found that contemporary privileged consumers seem to focus less on impressing others and more on the intrinsic pleasure obtained from goods, as well as family- and self-improvement (Postrel, 2008; Makkar & Yap, 2018). As such, rather than spending on luxury items such as shark fins as a tool of public status competition, Singapore’s consumers may be more inclined to invest in other services or experiences that can be converted into relationship-building, as well as lower visibility goods such as education, health care and retirement savings (Eckhardt, Belk & Wilson, 2015; Currid-Halkett, 2017). Moreover, the importance of modesty and fear of being targets of resentment and envy instilled in the rich in Singapore may have prompted consumers to avoid forms of conspicuous consumption which overtly display their wealth (Makkar & Yap, 2018; Ng, 2021; Tan, 2021; Mokhtar, 2022; Shehnaz, 2022).

It is also noteworthy that the minority of respondents who preferred higher quality fins also had the strongest preference for lower-priced fins (LCM Segment 2). While a ‘high quality, low price’ preference may seem a contradiction due to the mismatch in cues, it has been shown to motivate purchase intentions when consumers have weak price-quality schema (i.e., do not consider the purchase of high-priced products to acquire high quality products) and low need for cognition (i.e., form judgements based on simple peripheral cues inherent in the advertisements) (Shirai, 2015). Previous studies also indicate that ‘high quality, low price’ appeals to consumers in Singapore, as evidenced by readiness to queue for a good deal and desire to maximize the value of each dollar spent with good quality (Setiawan, 2020). This concern with getting the most out of sales or promotions likely stems from the *kiasu* (the fear of losing out or being left behind) mindset in Singapore (Bedford & Chua, 2018). Moreover, when faced with inconsistent cues simultaneously, consumers in Singapore are likely to value ‘low price’ over ‘high quality’ for high-priced products such as shark fins (Shirai, 2015). Consumers in LCM Segment 2 were also less likely to accept large price premiums for alternatives as compared to other segments (e.g., LCM Segment 2 was only willing to accept SGD $0.60 for lab-cultured shark fins versus SGD $3.56 for LCM Segment 1). This segment is therefore characterised by price-conscious consumers, who focus on low prices and have narrower latitudes of acceptable prices (McDaniel, Rao & Jackson, 1986; Shirai, 2015).

#### 4.1.2 Preferences for responsibly-sourced and lab-cultured shark fins over wild-caught shark fins from unmanaged sources

Consumers across all segments exhibited preferences for responsibly sourced wild-caught shark fins over wild-caught shark fins from unmanaged sources. These findings corroborate with a previous study which reported reduced shark fin consumption due to shark conservation and environmental concerns, as well as willingness to support restaurants that serve responsibly sourced and sustainable seafood (WWF, 2016). Indeed, consumers in Singapore are becoming increasingly aware and supportive of sustainable seafood (Jones, 2015; Gabriel, 2020).

Similarly, more than 80% of respondents preferred lab-cultured shark fins over unmanaged wild-sourced shark fins (LCM Segment 1, 2, 4). This receptivity to lab-cultured alternatives is possibly again due to the Singapore cultural trait of *kiasuism*, which may motivate consumers to project an image of being ‘ahead of the curve’ in their thinking and behavior (Chong, Leung & Lua, 2022). Specifically, a small group of consumers (LCM Segment 4) was willing to pay approximately 50% more for lab-cultured shark fins than for responsibly wild-sourced shark fins, suggesting that lab-cultured shark fins could find a high value market niche among early-adopters. Although our study did not find any demographic characteristics associated with LCM Segment 4, several studies have observed that potential consumers of cultured meat are young, highly educated, male, city dwelling, meat-consumers, and have a specific understanding of cultured meat (Tucker, 2014; Wilks & Phillips, 2017; Slade, 2018; Mancini & Antonioli, 2020; Zhang et al., 2021).

However, while our results showed that replacing unmanaged wild-sourced shark fins with lab-cultured shark fins could be a viable option to relieve harvest pressure on shark populations, two thirds of respondents (LCM Segment 1 and 3) had a higher willingness to pay for responsibly-sourced wild-caught shark fins over lab-cultured shark fins. Specifically, a minority of consumers (LCM Segment 3) strongly disliked lab-cultured fins and were unwilling to pay more. Prior research has suggested that some consumers generally oppose lab-cultured meat because of perceived unnaturalness, distrust of biotechnology, health concerns, taste, texture, and price (Hwang et al., 2020; Mancini & Antonioli, 2020; Zhang et al., 2021; Chriki, Ellies-Oury & Hocquette, 2022).

In addition, consumers in LCM Segment 3 had egoistically-based environmental concerns and do not view themselves as interconnected with other people or the natural environment (Schultz, 2000). This however does not mean that people with egoistic attitudes are unconcerned or apathetic about environmental problems, but rather they are more concerned about specific issues that directly impact them, their health and their future, and are thus motivated by reward or consequences for themselves (Schultz, 2000, 2001). Such orientation has implications for informing efforts to promote widespread collective environmental actions (as well as consuming lab-cultured shark fins), as these individuals would be more likely to engage in pro-environmental behaviors only when the action serves their personal needs and wants (Kollmuss & Agyeman, 2002; De Dominicis, Schultz & Bonaiuto, 2017).

#### 4.1.3 Passive consumption of shark fins

More broadly, shark fin sales to hotels and restaurants and prices have plummeted in Singapore since 2013, with retailers reporting recent inventory stockpiles (WWF, 2016; Boon, 2017; Choy et al., 2022). These changes in the shark fin market could be attributable in part to low and declining consumption of shark fins, and our study provides further empirical evidence to support this. Most respondents reported having only consumed shark fin a few times, once or less than once per year; and more than half of the sample reported decreased consumption within the past five years. Furthermore, respondents reported often consuming shark fins because it was served to them at weddings or special celebratory events. A small proportion of respondents (6.5%) also cited that fewer weddings and celebratory events due to the COVID pandemic has resulted in a decline in their shark fin consumption. This finding is supported by results of a consumer survey conducted by WWF in 2016, revealing that shark fin is most likely to be consumed at wedding banquets, and traditions and celebrations are the main reasons for consuming it (WWF, 2016). A great deal of consumption therefore appears to be passive rather than diners asking specifically for shark fin.

However, the continued inclusion of shark fin dishes in default set menus for banquets, Lunar New Year, or high-end restaurants does little to translate changing consumer sentiment into conservation outcomes (Sadovy de Mitcheson et al., 2018). Passive consumption of shark fins also has implications for how alternatives are priced in the market. While most respondents earned close to the average salary in Singapore (Koo, 2022), they do not typically actively consume shark fins, and thus are less inclined to find alternatives and pay a premium for these products. As such, it is key to consider the important role of passive consumption in any strategy to reduce consumption, suggesting that beyond consumers, it is also important to work with retailers and organizers of large events and celebrations to at least increase the agency of consumers regarding the shark fins that are served. It is unfortunate that passive consumers did not feature particularly in any of the segments in our model, as that makes it impossible to better characterize their demographics and psychographics.

On the other hand, a small group of respondents indicated either an increase or no change in consumption within the past five years and cited the dish’s delicious taste, nutritional value, and health benefits as motivating factors (Supplemental Material, Table S3). Indeed, shark fin soup has held perceived medicinal value for thousands of years, and adherents of traditional Chinese medicine believe that shark fins provide benefits as a tonic and have anti-cancer properties (Jarvis, 2019). However, contrary to these beliefs, shark fins can contain high levels of toxic mercury, which can lead to several adverse health risks when ingested (Choy & Wainwright, 2022). Therefore, improved culturally-relevant consumer education could help to alleviate misconceptions about the health benefits of shark fin. Furthermore, shark fins are essentially tasteless and the soup flavor comes from other ingredients used in the preparation of the broth (Dell’Apa, Chad Smith & Kaneshiro-Pineiro, 2014).

### 4.2 Methodological limitations

Discrete choice experiments (DCEs) have become an important tool for assessing consumer preferences for environmental goods, and can also elicit consumer preferences when goods and services are not yet offered or traded in the market (Rakotonarivo, Schaafsma & Hockley, 2016; Terris-Prestholt et al., 2019). As sustainable shark fins currently do not exist in the market (Zheng, 2017), we opted for an unlabeled DCE in this study, so that the alternatives in the choice set do not need to be described with labels such as brand names or logos. While these alternatives can be more abstract for respondents, unlabeled DCE’s have advantages such as (i) creating a design with attribute levels sufficiently broad to represent all possible alternatives, and (ii) encouraging respondents to trade off attribute levels of product that they are not familiar with for example because they are not yet widely available (de Bekker-Grob et al., 2010).

In addition, instead of investigating preferences of both consumers and non-consumers, we focused on recent shark fin consumers in the Singapore for our DCE. This criterion encompassed a wide range of demographics, with the distribution of race and age across our respondents representing the composition of the residential population in Singapore. We however found differences in other categories (e.g., gender, education level and monthly income) between our respondents and the resident population in Singapore. Future choice experiments can be refined with the intent to sample respondents representing the resident population so that DCE results are more likely to be reflective of actual domestic buyers, applicable to the Singapore population and relevant to future demand reduction efforts that can be implemented within the country (Doughty et al., 2021a). Additionally, we used latent class modelling to identify the preferences of the different consumer segments within sample population who shared similar outward patterns and characteristics (Weller, Bowen & Faubert, 2020). Taken together, these methodological decisions make this research more likely to yield actionable insights that can inform conservation efforts.

One limitation of our study is that we found passive consumers (who reported wedding and celebratory events to be the common setting for consuming shark fin) comprised 55% of our sample population, while our choice experiment was designed based on an active consumption scenario, which therefore still accounts for the most frequent context for nearly half of respondents which is non-negligible. Moreover, consumers who report passive consumption as the most frequent form of consumption may still actively consume shark fin in other contexts. In the future, our research could be supplemented by choice experiments which explore other target audiences that reflect a passive consumption scenario, such as recently married couples or caterers. Furthermore, passive consumption describes the guest’s state of consumption, but does not describe the host’s drivers and motivations for consumption at banquets and events. For instance, the cost of hosting a banquet may be split among the wedding couples’ families or solely by the parents hence a more senior family member could possibly decide to include shark fin in the menu (Teo, 2015; Singapore Brides, 2018). As such, future work could also focus on understanding who makes the ultimate decision of choosing shark fin at banquets and events, along with their motivations and drivers.

### 4.3 Leveraging preferences to deliver sustainability

Our research provides insights for three types of interventions to alleviate shark fishing pressure: (i) market-based interventions relating to sustainable fisheries and substitute lab-cultured products; (ii) government-led policy-level interventions on the sale and trade of shark products; and (iii) targeted demand-reduction efforts for different consumer segments to ensure market demand remains within the limits of sustainable supply. Drawing on a complementary mix of these strategies could help to reach the largest number of consumers (given heterogeneity in consumer segments) and thus have the greatest positive impacts.

#### 4.3.1 Market-based interventions

Our findings highlighted a strong market potential for sustainably sourced shark fins in Singapore, which could be fulfilled through two complementary avenues: limited sourcing from responsible fisheries, alongside supplementing demand with lab-cultured substitutes.

While the idea of responsibly-sourced shark fins is appealing, achieving a scalable market remains challenging. Most of the shark species currently managed for sustainability are not appropriate for the fin trade, and only 8.7% of fins in global fin trade are thought to originate from sustainable sources (Simpfendorfer & Dulvy, 2017; Zhou et al., 2021). These sustainably sourced fins are however not yet traceable or labeled, and do not stand out have enough visibility in the retail market to command price premiums. As such, it is difficult to leverage consumer preferences to incentivize more fisheries to adopt best-practice sustainability measures for more stocks (Simpfendorfer & Dulvy, 2017; Zhou et al., 2021).

In Singapore, some of these challenges could be overcome through direct purchasing or improved market integration between retailers and fishers. One option could involve specialized retailers which market responsibly-sourced shark fins to consumers, with price premiums paid to small-scale lower-impact fisheries operating in South East Asia, on the condition that certain sustainability standards are met, such as only capturing faster growing non-endangered species (e.g., blue sharks (*Prionace glauca*) and silky sharks (*Carcharhinus falciformis*), which are commonly found in the market currently, have growth rates fast enough to withstand some fishing pressure (Simpfendorfer & Dulvy, 2017; Zhou et al., 2021)); safely releasing endangered species (Booth et al., 2023); and ensuring key fisheries indicators (e.g., total catch, catch per unit effort and size-based indicators) are kept within science-based limits. This could not only help to incentivize better fishing practices in a conservation priority region, but also support the livelihoods of small-scale fishers (Booth, Squires & Milner-Gulland, 2019; Booth et al., 2021b).

Besides responsibly-sourced wild-caught shark fins, our results show that lab-cultured shark fins hold great promise to supplement the market and meet consumer demand. Such a product could be particularly popular in a collectivistic country like Singapore, where people are typically concerned with presenting a desirable impression of themselves or gaining higher prestige by using or endorsing a product that is visibly popular among others (Chong, Leung & Lua, 2022). Against this background, the generally positive attitude towards lab-cultured shark fins identified in our study indicates an opportunity for the alternative protein industry to prioritize product launches in Singapore, increasing the visibility of products using social media coverage. In addition, marketing strategies for lab-cultured shark fins could tap into the *kiasuism* mindset and ‘high quality, low price’ appeal by organizing sales and promotions along with detailed information on how low price was achieved while maintaining high quality (Shirai, 2015).

Yet despite their promise, some consumers (LCM Segment 3) remain skeptical towards lab-cultured shark fins. Future research could be conducted to examine the reasons associated with these consumer sentiments, to understand whether concerns surrounding sustainability and health benefits, perceived unnaturalness, or food technology neophobia are hindering their acceptance towards lab-cultured shark fins (Chong, Leung & Lua, 2022) is crucial to creating effective product messaging. Exposing skeptical consumers to lab-cultured shark fins through sensory tests, alongside messaging around the relative health benefits due to the lack of mercury contamination, could also perhaps increase consumers’ familiarity with products, and encourage them to choose it over fins sourced from unmanaged fisheries.

#### 4.3.2 Policy and regulatory interventions

To date, demand reduction campaigns in Singapore led by non-governmental organizations have largely engaged the food and beverage industry, with some large establishments and delivery services publicly committing to phase out shark fin (WWF, 2018). However, it appears that some restaurants may choose to serve shark fins again after the trial period, or upon request by regular customers (Yeo, 2022). Therefore, such self-regulation is voluntary and risks unfulfilled promises because of weak standards or ineffective enforcement, allowing establishments to continue to serve their own interests at the expense of public goods (Sharma, Teret & Brownell, 2010).

Given that many consumers in Singapore may consume shark fins passively, demand-reduction campaigns could also reach event management and wedding industries in addition to the food and beverage industry so that shark fins are excluded from their packages, or more sustainable options (such as responsibly-sourced or lab-grown) can be offered. Policies that aim to achieve industry-wide reform on the sale and trade of shark fins should also be considered. Indeed, the majority of consumers in Singapore believe that the government is not doing enough to protect sharks, and showed support for government legislation to restrict domestic consumption and trade in shark fins (WWF, 2016).

Possible policy options include: species-specific sales restrictions for Critically Endangered species such as wedgefish and hammerhead sharks; quota and permitting systems, which require traders and retailers to prove that purchasing and trade of sharks does not threaten wild populations (e.g., as per Appendix II of the Convention on the International Trade of Endangered Species); and taxes on shark fins, which may increase its price beyond consumers’ willingness to pay or prompt the supply chain to reduce the amount of shark fin imports because of lower profit from sales. Although taxes may not always be effective to shift behaviors in heterogeneous consumer markets, a preference towards lower-priced fins among consumers in Singapore suggest that demand could potentially be curbed this way, while taxes on seafood products could also raise government revenue for investment in marine conservation actions (Booth et al., 2021a). The Singapore government could set an example for consumers to reduce consumption by echoing China’s decision to cease serving it during official functions (Wassener, 2012; Zheng, 2019).

More broadly, since many shark and ray species migrate across international borders and are landed in multiple fisheries globally, international partnerships and collaboration across broad range of private and public sector institutions are necessary to deliver conservation success. As a global entrepot for shark fin trade, retailers and regulators in Singapore could develop multi-lateral collaborations with governments and fishers shark fishing countries in Southeast Asia – such as Indonesia and Malaysia – to promote improved systems of traceability from producer to consumer, which could enable responsibly-sourced shark fin to become a reality. Indeed, an integrated mix of management measures beyond consumer interventions and Singapore’s borders - such as science-based catch limits and protection of critical habitat – are needed to maintain the health of shark populations globally (Dulvy et al., 2014).

#### 4.3.3 Demand reduction

Finally, while local demand for shark fins appears to be shrinking in Singapore, further demand reduction initiatives could continue to reinforce consumers’ new beliefs, strengthen their commitment to behavioral shifts, and sustain lasting changes in consumption and preferences. While the importance of food safety was not tested due to limitations on the number of attributes that respondents can meaningfully trade-off, consumers in Singapore are increasingly embracing healthier diets and are motivated to adopt healthy lifestyles (Yeo, 2021). As such, campaigns dispelling myths relating to the health benefits of shark fins, and highlighting the potential negative impacts of elevated levels of mercury, could be effective. This may be particularly impactful for those in LCM Segment 3 who placed more value on their future, health, and lifestyle, and could therefore motivate them to change their consumption behaviors (De Dominicis, Schultz & Bonaiuto, 2017). Messaging focused on self-interest could also be complemented with messaging on altruistic values (De Dominicis, Schultz & Bonaiuto, 2017), such as shark endangerment, which could be most relevant to consumers in LCM Segment 1 and 2 whose environmental concerns were less oriented around self-interest and more related to ecosystems and others. Consumers in LCM Segment 2 who values fin quality could also be receptive to messages focusing on counterfeit shark fins (Zhou et al., 2021).

Since many respondents reported passive consumptions, more work needs to be done to understand, for example, how menus are designed for events and banquets, who are the decision makers in this context, and how they can be influenced by strong social and cultural norms. It is possible that venues, event planners and caterers, as well as close relatives, could play an important role in these decisions (Lo et al., 2021), which could make them critical audiences for future demand reduction interventions.

### 4.4 Wider implications for humans and nature

While this study focuses specifically on shark fin consumption in the context of Singapore, the findings reflect similar research from other cases of consumption of endangered species in urban areas for non-subsistence purposes. For example, social context and strongly held (but not scientifically proven) beliefs also drive consumption of pangolins in Ho Chi Min city, saiga horn in Singapore and wild game meat in Poland (Doughty et al., 2019, 2021a; Olmedo et al., 2022). This highlights the complexity of demand for wildlife products, and the importance of understanding the social meaning of wildlife products to propose effective solutions. Moreover, this is not only relevant for changing consumer behaviour, but applies to all forms of interventions which seek to change human behavior for conservation purposes. There is a need to understand the economic and social drivers of behaviors which lead to the overexploitation of sharks and other endangered wildlife to develop solutions that are both effective and socially equitable (John et al., 2010; Veríssimo, 2013; Booth, Squires & Milner-Gulland, 2019).

## 5. Concluding remarks

A comprehensive understanding of consumer behaviors and preferences is a crucial first step to designing effective interventions that stem demand for wildlife products (Doughty et al., 2021b). Yet despite being a global priority country for shark management, very little data exists on consumer preferences for shark fins in Singapore. Our study thus provides much-needed evidence on the characteristics and preferences of shark fin consumers in Singapore, which can be leveraged to design future interventions to curb the unsustainable use of shark fins. Our results highlight that shark fins sourced from responsible fisheries and lab-cultured alternatives offer promising substitutes to those sourced from unmanaged fisheries. Based on these findings, we call on the government, businesses, and consumers in Singapore to take the lead in testing novel market-based and policy solutions, which could create positive rippling effects on global shark and ray populations and coastal communities.

## CRediT authorship contribution statement

Christina Choy (CC), Hollie Booth (HB) and Diogo Veríssimo (DV): Conceptualization, Methodology, Investigation, Data curation, Formal analysis, Writing - Original Draft, Writing - Review & Editing, Project administration, Funding acquisition

## Data availability

The data that support the findings of this study will be openly available in Zenodo or Harvard Dataverse.

## Acknowledgements

This research was possible thanks to the funding offered by Silverstrand Capital to the Wildlife Conservation Society, and their unwavering support. All authors would like to thank our friends and colleagues in Singapore who have helped with the survey pilot-test, offered their feedback, and enriched our research. We also thank the anonymous reviewers whose comments greatly helped in improving the quality of the manuscript.

## List of Abbreviations

AIC: Akaike Information Criteria
ASC: Alternative Specific Constant
BIC: Bayesian Information Criteria
DCE: Discrete Choice Experiment
LCM: Latent Class Model
MNL: Multinomial Logit
SGD: Singapore Dollars

