## Supplemental Material for "Understanding consumers to inform market interventions for Singapore’s shark fin trade"

Survey questionnaire

Part I - Demographic information

Q1. What is your age?
- a. 21-25
- b. 26-30
- c. 31-35
- d. 36-40
- e. 41-45
- f. 46-50
- g. 51-55
- h. 56-60
- i. Above 60

Q2. To which gender identity do you most identify?
- a. Male
- b. Female
- c. Gender variant/non-conforming
- d. Prefer not to answer

Q3. What is your race?
- a. Chinese
- b. Malay
- c. Indian
- d. Others: __________ (please specify)

Q3a. If you are Chinese, which dialect group are you from?
- a. Cantonese
- b. Hakka
- c. Hokkien
- d. Peranakan
- e. Teochew
- f. Others: __________ (please specify)

Q4. What is your religion?
- a. Buddhism
- b. Christanity
- c. Hinduism
- d. Islam
- e. Roman Catholicism
- f. Taoism
- g. Others
- h. No religion

Q5. What is your highest education level?
- a. Below secondary
- b. Secondary
- c. Post-secondary (non-tertiary)
- d. Diploma and professional qualification
- e. University
- f. Post-graduate

Q6. What is your monthly income range (in Singapore Dollars)?

a. Below $1,000
b. $1,000 - $2,999
c. $3,000 - $4,999
d. $5,000 - $6,999
e. $7,000 - $8,999
f. $9,000 - $10,999
g. $11,000 - $12,999
h. $13,000 - $14,999
i. $15,000 - $17,499
j. $17,500 - $19,999
k. $20,000 and above

<u>Part II - Purchase and consumption habits</u>

Q7. When was the last time you consumed shark fin in Singapore?
a. Within the past week
b. Within the past month
c. Within the past six months
d. Within the past year

Q8. How frequently do you consume shark fin in Singapore?
a. Less than once a year
b. Once a year
c. A few times a year
d. At least once per month
e. At least once per week

Q9. In which of the following situations do you most frequently consume shark fin in Singapore? Please select one.
a. Business meetings
b. Weddings or special celebratory events
c. Dine-out meals with family and friends
d. At home
e. Others: __________ (please specify)

Q10. Which of the following best describes your shark fin consumption in Singapore?
a. I purchase dried or instant frozen shark fins directly from a supplier
b. I purchase cooked shark fin soup in restaurants
c. I don’t purchase it myself, but I eat it at a wedding or special celebratory event if it is provided by someone else
d. Others: __________ (please specify)

Q11. Where have you purchased shark fin (dried, instant frozen or chilled, cooked soup) in Singapore in the past? Please select all that apply.
a. Restaurants or hotels
b. Fishery ports
c. Wet markets (e.g., Tekka Market, Chinatown Market)
d. Supermarkets (e.g. NTUC, Sheng Siong)
e. Dried seafood shops
f. Traditional medicinal halls
g. Online
h. I do not purchase shark fin
i. Others: __________ (please specify)

Q12. Has your shark fin consumption changed in the past 5 years?
a. Large decrease
b. Small decrease
c. No change
d. Small increase
e. Large increase

Q13. Please explain where applicable, why did you change or not change your shark fin consumption?

_______________________________________________________________________

_______________________________________________________________________

_______________________________________________________________________

________

Part III - Pro environmental attitudes

Q14. To what extent do you agree/disagree with the following statements:

| | **Strongly disagree** | **Disagree** | **Neither disagree or agree** | **Agree** | **Strongly agree** |
|---|---|---|---|---|---|
| a. *I like to eat shark fin because it is delicious* | | | | | |
| b. *I like to eat shark fin because it is prestigious* | | | | | |
| *c. I like to eat shark fin because it is traditional* | | | | | |
| *d. I like to eat shark fin because its sociable* | | | | | |
| *e. I like to eat shark fin because its nutritious* | | | | | |
| *f. We should manage shark populations to sustain other fish stocks* | | | | | |
| *g. Sharks are important for the functioning of marine ecosystems* | | | | | |

Q15. People around the world are generally concerned about environmental problems because of the consequences that result from harming nature. However, people differ in the consequences that concern them the most.

Please rate each of the following items from 1 (Not important) to 7 (Supreme importance) in response to the question:

*I am concerned about environmental problems because of the consequences for*_____________.

Plants
Marine life
Birds
Animals
Me

My health
My future
All people
Children

<u>Part IV - Choice Experiment</u>

Q16. Imagine you are dining out with your family (with older folks e.g, parents and grandparents) at a Chinese restaurant which serves shark's fin with crab roe. There are three options on the menu for shark fin soup as follows; which choice would you prefer?

Each choice below varies in terms of the following five attributes: price per person serving, size of shark fin, texture and quality of shark fin, menu types for placing an order, and information on source of shark fin. Please indicate which choice you would be likely to choose if offered to you in real life. You will be asked to make a total of 15 choices regarding your preference for different shark fin soups, and the choices are randomized. There are no right or wrong answers. We are simply interested to understand the preferences of Singaporean shark fin consumers.

For example,

| (a) | (b) | (c) | (d) Another dish that does not have shark fin |
|---|---|---|---|
| 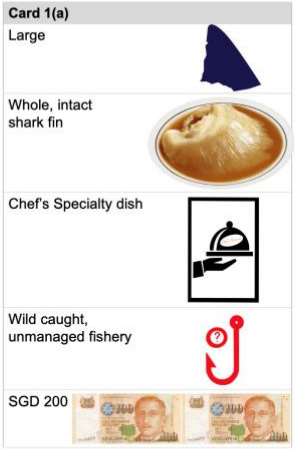 | 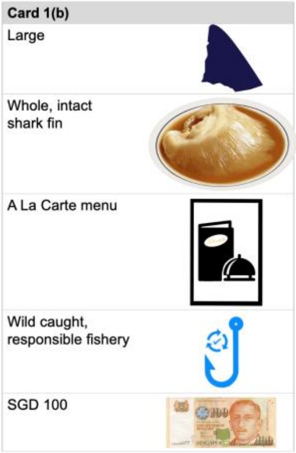 | 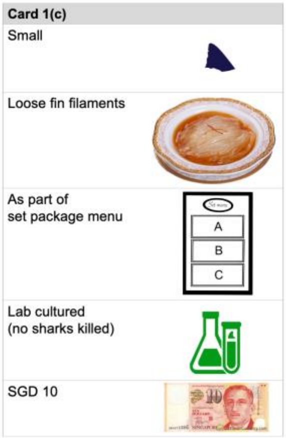 | |

Combinations used in the survey questionnaire

Legend

| Attribute | Level | Code |
|---|---|---|
| Source | Wild caught, unmanaged fishery | 0 |
| | Wild caught, responsible fishery | 1 |
| | Lab cultured (no sharks killed) | 2 |
| Menu | As part of set package menu | 0 |
| | A La Carte menu | 1 |
| | Chef's Specialty dish | 2 |
| Quality | Loose fin filaments | 0 |
| | Whole, intact shark fin | 1 |
| Size | Small | 0 |
| | Large | 1 |
| Price | SGD 10 | 0 |
| | SGD 50 | 1 |
| | SGD100 | 2 |
| | SGD150 | 3 |
| | SGD 200 | 4 |

| Option | Attribute code | | | | |
|---|---|---|---|---|---|
| | Source | Menu | Quality | Size | Price |
| 1a | 0 | 2 | 1 | 1 | 4 |
| 1b | 1 | 1 | 1 | 1 | 2 |
| 1c | 2 | 0 | 0 | 0 | 0 |
| 2a | 0 | 0 | 1 | 1 | 1 |
| 2b | 2 | 2 | 0 | 0 | 4 |
| 2c | 1 | 1 | 1 | 1 | 2 |
| 3a | 2 | 2 | 1 | 1 | 0 |
| 3b | 1 | 1 | 0 | 0 | 2 |
| 3c | 0 | 0 | 0 | 0 | 4 |
| 4a | 1 | 1 | 0 | 0 | 3 |
| 4b | 2 | 0 | 1 | 1 | 0 |
| 4c | 0 | 2 | 0 | 0 | 3 |
| 5a | 1 | 1 | 0 | 0 | 2 |
| 5b | 0 | 2 | 0 | 1 | 0 |
| 5c | 2 | 0 | 1 | 0 | 4 |
| 6a | 0 | 2 | 1 | 0 | 0 |
| 6b | 2 | 0 | 0 | 1 | 4 |
| 6c | 1 | 1 | 1 | 0 | 2 |
| 7a | 1 | 1 | 0 | 0 | 2 |
| 7b | 0 | 0 | 1 | 0 | 3 |
| 7c | 2 | 2 | 0 | 1 | 1 |
| 8a | 0 | 2 | 0 | 1 | 4 |
| 8b | 1 | 1 | 0 | 1 | 2 |
| 8c | 2 | 0 | 1 | 0 | 0 |
| 9a | 2 | 0 | 0 | 1 | 0 |
| 9b | 1 | 1 | 1 | 0 | 3 |

| | | | | | |
|---|---|---|---|---|---|
| 9c | 0 | 2 | 1 | 0 | 3 |
| 10a | 0 | 0 | 1 | 1 | 3 |
| 10b | 2 | 2 | 0 | 0 | 0 |
| 10c | 1 | 1 | 1 | 1 | 3 |
| 11a | 1 | 1 | 0 | 0 | 2 |
| 11b | 2 | 2 | 1 | 0 | 4 |
| 11c | 0 | 0 | 0 | 1 | 0 |
| 12a | 2 | 2 | 1 | 0 | 1 |
| 12b | 0 | 0 | 0 | 1 | 3 |
| 12c | 1 | 1 | 0 | 1 | 2 |
| 13a | 1 | 1 | 0 | 0 | 1 |
| 13b | 0 | 0 | 1 | 0 | 1 |
| 13c | 2 | 2 | 0 | 1 | 4 |
| 14a | 2 | 0 | 0 | 0 | 3 |
| 14b | 1 | 1 | 1 | 1 | 1 |
| 14c | 0 | 2 | 1 | 1 | 1 |
| 15a | 2 | 0 | 1 | 1 | 4 |
| 15b | 0 | 2 | 0 | 0 | 1 |
| 15c | 1 | 1 | 0 | 0 | 1 |

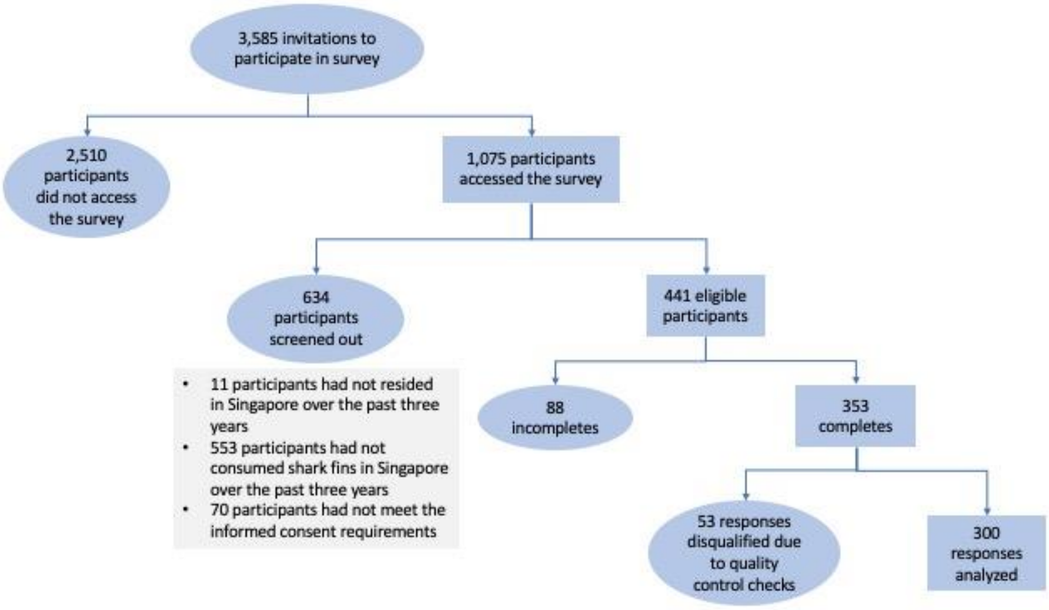


Figure S1. Summary of the survey dispositions

Table S2. Summary of respondent demographics

| Variable | Description | Response (%) |
|---|---|---|
| Age | 21-30 | 17 |
| | 31-40 | 18 |
| | 41-50 | 22 |
| | 51-60 | 22 |
| | 61 and above | 22 |
| Gender | Male | 60 |
| | Female | 40 |
| Race | Chinese | 76 |
| | Malay | 14 |
| | Indian | 7 |
| | Others | 3 |
| Dialect group | Cantonese | 28 |
| | Hakka | 7 |
| | Hokkien | 38 |
| | Peranakan | 2 |
| | Teochew | 22 |
| | Others | 3 |
| Religion | Buddhism | 25 |
| | Christanity | 33 |
| | Hinduism | 4 |
| | Islam | 9 |
| | Roman Catholicism | 7 |
| | Taoism | 7 |
| | Others | 1 |
| | No religion | 15 |
| Education level | Below secondary | 0 |
| | Secondary | 6 |
| | Post-secondary (non-tertiary) | 6 |
| | Diploma and professional qualifications | 17 |

| | | |
|---|---|---|
| | University | 51 |
| | Post-graduate | 20 |
| Monthly income range | Below $3,000 | 14 |
| | $3,000 - $7,000 | 40 |
| | $7,000 - $11,000 | 22 |
| | $11,000 - $15,000 | 11 |
| | $15,000 and above | 13 |

Table S3. Summary of respondents' purchase and consumption habits

| Question | Description | Response (%) |
|---|---|---|
| When was the last time you consumed shark fin in Singapore? | Within the past week | 20 |
| | Within the past month | 25 |
| | Within the past six months | 22 |
| | Within the past year | 33 |
| How frequently do you consume shark fin in Singapore? | Less than once a year | 17 |
| | Once a year | 19 |
| | A few times a year | 36 |
| | At least once per month | 13 |
| | At least once per week | 15 |
| In which of the following situations do you most frequently consume shark fin in Singapore? | Business meetings | 10 |
| | Weddings or special celebratory events | 55 |
| | Dine-out meals with family and friends | 20 |
| | At home | 15 |
| Which of the following best describes your shark fin consumption in Singapore? | I purchase dried or instant frozen shark fins directly from a supplier | 23 |
| | I purchase cooked shark fin soup in restaurants | 24 |
| | I don't purchase it myself, but I eat it at a wedding or special celebratory event if it is provided by someone | 52 |
| | Others | 1 |
| Has your shark fin consumption changed in the past 5 years | Large decrease | 30 |
| | Small decrease | 31 |
| | No change | 26 |
| | Small increase | 6 |
| | Large increase | 7 |
| Reasons for increased consumption within the past 5 years | ● I love to consume it and happy to use it<br>● Sharks fin is a healthy food and has sufficient supply anywhere in the world<br>● My personal preference<br>● It tastes good each day<br>● More healthy<br>● I realized the nutritious value of shark fin<br>● They are delicious and nutritious too<br>● Related to me | |

- I love the taste of sharks fin. Braised or stewed or with crab meat soup is the best. It's a luxury food fit for wealthy millionaires like myself.
- Because i have grown a lot of interest in it
- I increased my consumption because my family members like it
- I found more passion and my family too developed a great taste
- Because I like to eat Shark fins
- It is good service. So i do this.
- Before shark little bit places available but now many places outlets is good consumption
- Nutritious
- They're more delicious and nutritious too
- Delicious, nutritious and sociable
- Very good
- They're delicious and nutritious
- Related to my requirements
- It is good
- I increased my consumption because I read about how nutritional it is
- Nice
- They're delicious, prestigious, nutritious and sociable
- My favorites shark fin taste good and delicious
- I sticked to what i eat and won't change to any
- Very tasty and healthy
- Which is healthy and tasty
- Good for health
- I found them being delicious
- It is not widely available like before
- Shark fin is expensive products.
- Good in collagen and have class eating shark fin Business partner treat me, I treat them
- Its a special occasion and used for celebratory period
- In recent times my consumption of shark fin has increased a little due to the fact that I now dine in more restaurants compared to previous years.

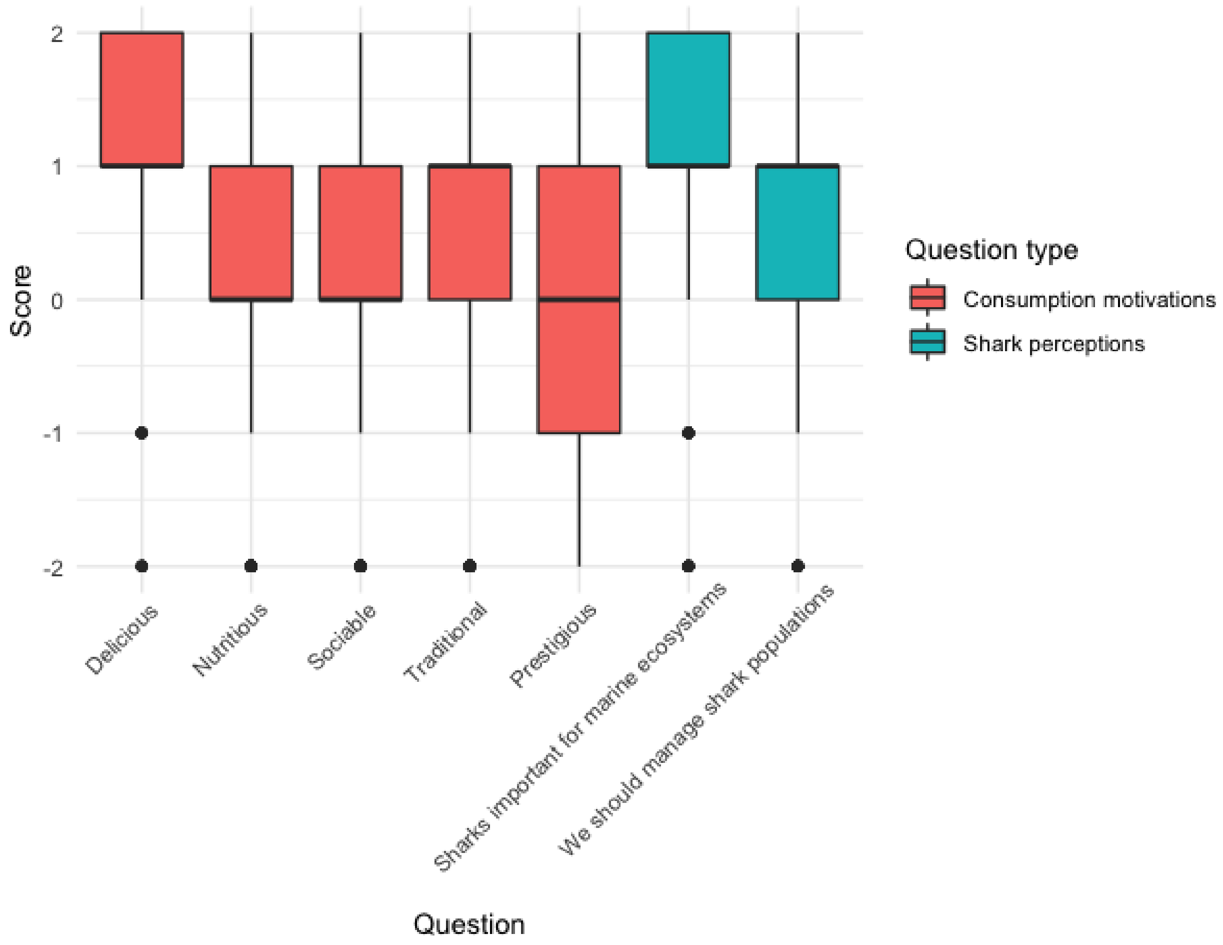


Figure S4. Mean attitude scores of respondents on consumption motivations and perceptions of sharks, with 95% confidence intervals. 1 = strongly disagree, 2 = disagree, 3 = neither disagree or agree, 4 = agree, 5 = strongly agree.

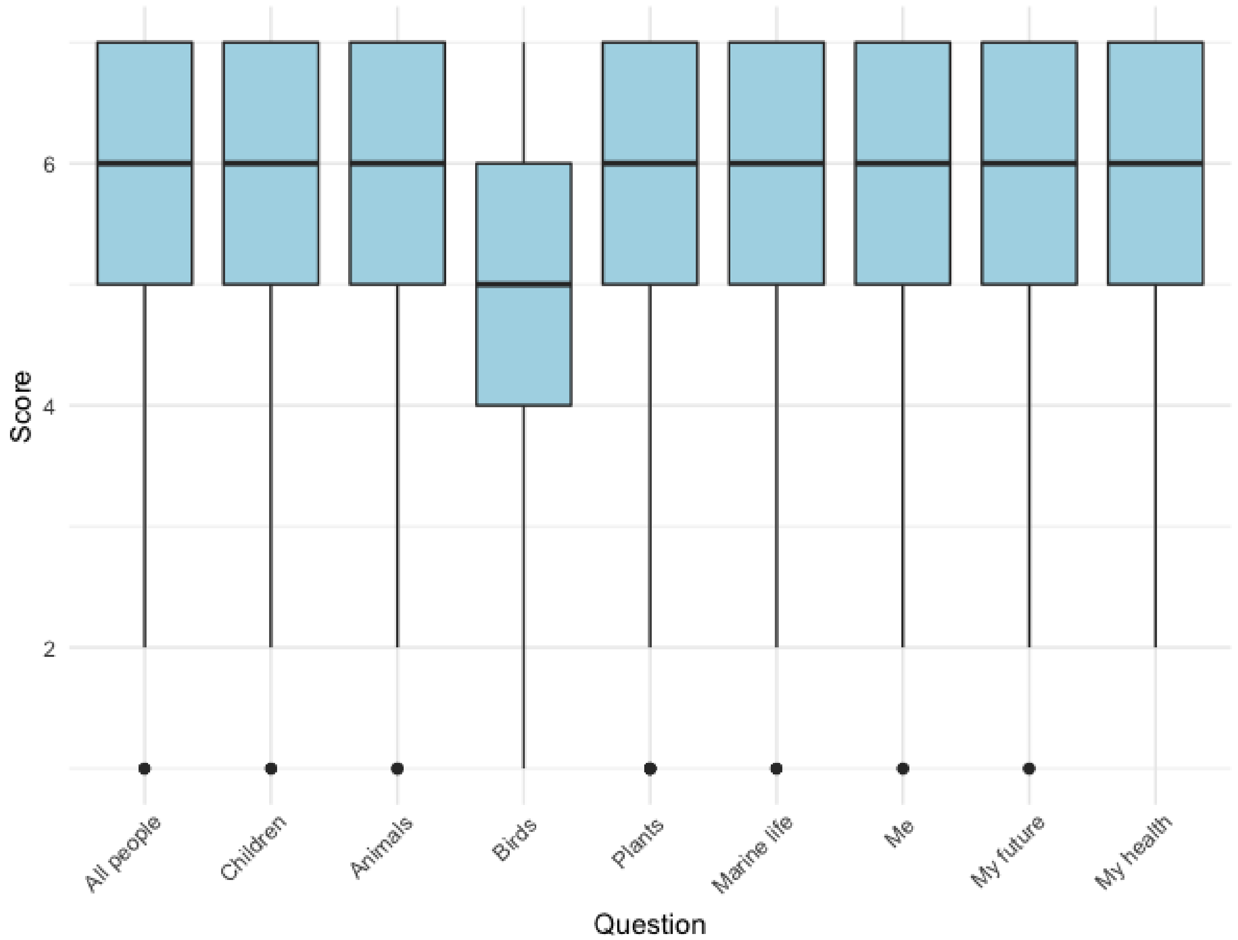


Figure S5. Mean attitude scores of respondents on environmental concern, with 95% confidence intervals, on a scale of 1 (not important) to 7 (supreme importance).

Table S6. Summary of measures of model fit for Multinomial Logit (MNL) and Latent Class Models (LCM). The selected model with best performance is underlined.

| Model | Number of parameters | Log Likelihood | Akaike's Information Criterion | Bayesian Information Criterion | AIC3[a] |
|---|---|---|---|---|---|
| MNL | 7 | -5654.455 | 11322.9 | | |
| 2 segment LCM | 19 | -4771.7 | 9581.5 | 9703 | 9600 |
| 3 segment LCM | 31 | -4518.5 | 9099 | 9298 | 9130 |
| <u>4 segment LCM</u> | <u>43</u> | <u>-4213.2</u> | <u>8512.4</u> | <u>8788</u> | <u>8555</u> |
| 5 segment LCM | 55 | -4149.2 | 8408.4 | 8761 | 8463 |

[a] Modified Akaike's Information Criterion with 3 as penalty factor
